# μCB-seq: Microfluidic cell barcoding and sequencing for high-resolution imaging and sequencing of single cells

**DOI:** 10.1101/2020.02.18.954974

**Authors:** Tyler N. Chen, Anushka Gupta, Mansi Zalavadia, Aaron M. Streets

**Affiliations:** University of California, Berkeley, Department of Bioengineering, Berkeley, CA; University of California, Berkeley, Department of Materials Science and Engineering, Berkeley, CA; UC Berkeley-UC San Francisco Graduate Program in Bioengineering, Berkeley, CA; Chan Zuckerberg Biohub, San Francisco, CA

## Abstract

Single-cell RNA sequencing (scRNA-seq) enables the investigation of complex biological processes in multicellular organisms with high resolution. However, many phenotypic features that are critical to understanding the functional role of cells in a heterogeneous tissue or organ are not directly encoded in the genome and therefore cannot be profiled with scRNA-seq. Quantitative optical microscopy has long been a powerful approach for characterizing diverse cellular phenotypes including cell morphology, protein localization, and chemical composition. Combining scRNA-seq with optical imaging has the potential to provide comprehensive single-cell analysis, allowing for functional integration of gene expression profiling and cell-state characterization. However, it is difficult to track single cells through both measurements; therefore, coupling current scRNA-seq protocols with optical measurements remains a challenge. Here, we report Microfluidic Cell Barcoding and Sequencing (μCB-seq), a microfluidic platform that combines high-resolution imaging and sequencing of single cells. *μ*CB-seq is enabled by a novel fabrication method that preloads primers with known barcode sequences inside addressable reaction chambers of a microfluidic device. In addition to enabling multi-modal single-cell analysis, μCB-seq improves gene detection sensitivity, providing a scalable and accurate method for information-rich characterization of single cells.

## Introduction

Over the last decade, single-cell genomics has revolutionized the study of complex biological systems,^1^ allowing us to map the composition of tissue,^2^ organs,^3-6^ and even whole organisms^7-9^ with unprecedented resolution. Most notably, single-cell RNA sequencing (scRNA-seq), which involves reverse transcription of mRNA followed by high-throughput sequencing of cDNA, has recently emerged as the most quantitative and comprehensive tool for profiling cellular identity. This technological revolution has been facilitated by the development of microfluidic workflows for scRNA-seq that make it possible to analyze hundreds to thousands of single cells in one experiment,^10-16^ paving the way for the construction of a human cell atlas.^17^ While scRNA-seq is effective for quantitatively measuring mRNA in large numbers of single cells, cellular identity is not entirely described by the transcriptome alone. Phenotypic features such as morphology, protein localization, and metabolic composition provide critical information about the identity, function, or state of cells but are not directly encoded in the genome, and therefore cannot be measured by sequencing. Thus, optical microscopy remains an indispensable tool for characterizing phenotypic features of single cells and multicellular systems. Combining microscopy with scRNA-seq can provide valuable insights into the relationship between gene expression and cellular phenotype. Furthermore, because each technique probes distinct features, imaging and sequencing single cells might provide a more comprehensive description of cellular identity.

Performing optical imaging and sequencing measurements on the same single cell is technically challenging because it requires precise cell manipulation and tracking. Microfluidic technology is well-suited to address such technical challenges, as it provides low Reynolds number, laminar flow, and programmable fluidic control at the microscale. Specifically, multi-layer microfluidic devices with integrated valves allow for the trapping and imaging of single cells which can then be sorted for downstream genomic analysis. For example, Lane *et al.* used microfluidic scRNA-seq with optical microscopy to combine fluorescent measurements of transcription factor dynamics with gene expression profiling in single cells.^18^ In their study, the link between a cell’s image and its transcriptome was preserved by carrying out individual library preparation for each cell, making library preparation the rate limiting step for total cell throughput. When imaging is not required, higher-throughput methods such as microwell- and microdroplet-based techniques allow for multiplexed processing of many cells at once, thus drastically reducing library preparation time.^11-16^ These methods use microfabricated devices to isolate cells in nanoliter volumes, in which cellular barcodes are incorporated into cDNA during reverse transcription (RT) to allow for pooling of many cells into a single sequencing library. However, these techniques are currently not compatible with imaging because cellular barcodes are assigned randomly, making it impossible to know which transcriptome belongs to which cell image. Yuan *et al.* recently demonstrated a promising solution to this challenge, in which the random barcode sequences were optically decoded using fluorescence microscopy.^19^ Spectrally-encoded beads^20^ or printed droplet microfluidics^21^ may provide yet other solutions for imaging and sequencing single cells. Despite these advances, further developments are needed to realize the benefits of combined high-resolution imaging and high-sensitivity RNA-seq on single cells.

In this report, we present Microfluidic Cell Barcoding and Sequencing (μCB-seq), a microfluidic platform that enables paired imaging and sequencing measurements of single cells. Our platform uses integrated microfluidic valves to precisely manipulate single cells for isolation, imaging, and multistep library preparation on-chip. In μCB-seq, independently addressable microfluidic reaction chambers are preloaded with known barcoded primers, which are used to capture genomic material from single cells. This approach provides the ability to couple genomic information with phenotypic information that requires high-resolution imaging or even time-resolved imaging to investigate dynamic cellular behaviour. Here, we demonstrate the capabilities of μCB-seq by performing scRNA-seq using the molecular crowding single-cell RNA barcoding and sequencing (mcSCRB-seq) protocol.^22^ We find that μCB-seq improves upon the high sensitivity of mcSCRB-seq by utilizing the benefits of microscale volume library preparation reactions.^23^ We then combine multiplexed scRNA-seq with live-cell fluorescence imaging on-chip to demonstrate μCB-seq as a scalable platform for extracting high-resolution phenotypic data and high-sensitivity genomic data from single cells.

## Results and Discussion

### 1. Microfluidic device design and μCB-seq workflow

μCB-seq is implemented on a PDMS microfluidic device with integrated elastomeric valves fabricated by multilayer soft-lithography.^24^ The device has two functional layers, an upper control layer, and a lower flow layer (Fig. 1A). The control valves are pneumatically actuated by a solenoid valve array that is operated with the KATARA controller and a programmable computer interface.^25^ The device design was inspired by a previous scRNA-seq platform,^23^ and in this demonstration, can process 10 cells simultaneously in parallel reaction lanes. Each reaction lane has a modular design to allow for imaging, cell lysis, and implementation of a wide range of multistep library preparation protocols. The imaging module consists of an imaging chamber flanked by two isolation valves (Fig. 1B), and the lysis and reverse transcription (RT) modules consist of isolated reaction chambers separated by valves (Fig. 1C, Fig. S1). During chip operation, a suspension of single cells is loaded into the cell inlet and directed towards the imaging module using pressure-driven flow. Once a cell reaches an imaging chamber, it is actively trapped, imaged, and then sorted into its respective reaction lane or sent to waste, allowing for the enrichment of cell subpopulations or the selection of rare cells. After imaging, the selected cell is ejected from the imaging chamber into the lysis module of its reaction lane by a flow of lysis buffer from the reagent inlet. After all 10 lysis modules are filled with lysis buffer, processing proceeds in parallel for all 10 cells.

**Figure 1:**
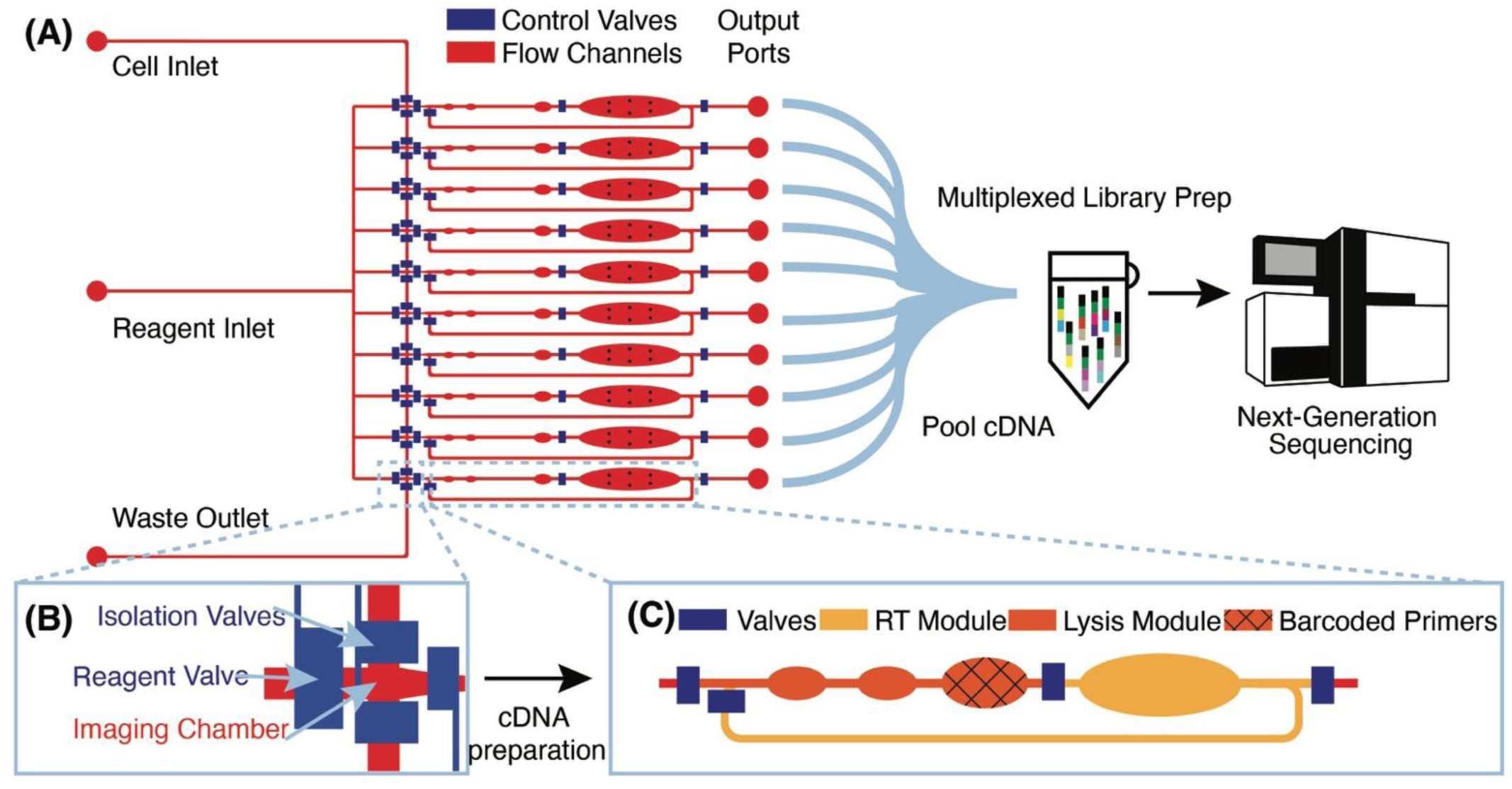
μCB-seq device design and workflow. (A) Schematic of the microfluidic device with control valves in blue and flow layer in red. Cells are loaded into the cell inlet and reagent is introduced through the reagent inlet. The device processes 10 cells in 10 individual reaction lanes, each ending in an output port. Reverse-transcribed cDNA is recovered from output ports for all cells, pooled in a single tube for off-chip library preparation using the mcSCRB-seq protocol, and sequenced using next-generation sequencing platforms. (B) Detailed diagram of the imaging module showing the imaging chamber. The two isolation valves can be actuated to actively capture a cell of interest in the imaging chamber. (C) Detailed diagram of one reaction lane showing the lysis and RT modules separated by valves. The textured reaction chamber in the lysis module is preloaded with barcoded RT primers.

RT primers with known barcode sequences are preloaded in the lysis module of each reaction lane (Fig. 1C, *Device Fabrication*). Each reaction lane is indexed by two pieces of information: a known barcode sequence and its lane index on the device. As a result, all sequencing reads with a unique cell barcode sequence can be linked to cell images with the corresponding lane index. Barcode sequences used in this study are a subset of 8-nt long Hamming-correctable barcodes^26^ designed for 50% GC content and minimal sequence redundancy (Table S1). The unique molecular identifier (UMI) sequence in the RT primers is 10-nt long.

Positioned above the reaction chambers in the lysis module are mixing paddles (Fig. S1), which are used to accelerate mixing as demonstrated previously.^23^ After dead-end filling of the lysis module, barcoded RT primers are resuspended in cell lysate by active mixing, after which the entire chip is placed on a temperature-controlled platform to hybridize suspended RT primers to cellular mRNA transcripts. The reagent input line is then flushed and filled with RT buffer, which is injected into all reaction lanes to dead-end fill the RT module. The RT buffer contains 7.5% PEG 8000, which has been demonstrated to increase RT efficiency through molecular crowding.^27,28^ Reverse transcription is carried out for 1.5 hours at 42°C, during which the mixing paddles are actuated in a peristaltic manner to circulate the relatively viscous RT mix throughout the mixing channel of each reaction lane (Fig. S1).

The total reaction volume of each lane is 227 nL, which is 1-2 orders of magnitude smaller than typical plate-based protocols.^22^ After RT, all lanes are independently flushed with 1.7 µL of nuclease-free water to recover cDNA, and pooled into a single tube using gel-loading pipette tips for a total volume of 17 μL. Additional off-chip steps including exonuclease digestion and cDNA amplification followed by purification and Nextera library preparation are performed in a single tube using the conventional mcSCRB-seq protocol *(Materials and Methods).* cDNA libraries representing whole single-cell transcriptomes are then sequenced on a next-generation sequencing platform.

### 2. Microfluidic device fabrication with addressable barcode spotting

Multilayer chip fabrication is necessary to create microfluidic devices with integrated valves and pumps that can be actuated for precise fluidic manipulation of cells, buffer exchange, and continuous-flow mixing of reagents.^29^ These capabilities enable the implementation of multistep reactions for library preparation on such devices. However, as the number of inputs and outputs are increased, “world-to-chip” interfacing becomes more complex.^30^ In order to increase multiplexing throughput while minimizing the complexity of device operation, μCB-seq utilizes a fabrication method that combines multilayer soft lithography and DNA array printing to preload the lysis module of each lane with known barcoded RT primers. This approach is similar to previous microfluidic devices for high-throughput screening of protein-DNA interactions.^31^ To verify that RT primers can be successfully resuspended from PDMS after baking, 2 μL droplets of 2 ng/μL primer were manually spotted on PDMS slabs, allowed to dry, baked at 80 °C for 2 hr, and incubated at room temperature for 24 hr. Primers were manually resuspended in 2 μL of nuclease-free water and analyzed for fragment length. The RT primers showed no noticeable degradation during the final baking at 80 °C and can be resuspended with high efficiency (Fig. S2).

The μCB-seq device was designed in the push-down configuration with three layers: a thick upper control layer, a thin middle flow layer, and a thin lower dummy layer. We used on-ratio PDMS-PDMS bonding to avoid PDMS waste and provide a stable seal by partial crosslinking of a 10:1 base:crosslinker mixture with each new layer of the microfluidic device.^32^ The control and flow molds were first patterned using standard photolithography techniques (Fig. 2A, *Materials and Methods)*. The 10:1 PDMS mixture was then separately cast onto the two molds and baked. The partially crosslinked control layer was peeled from the mold and placed atop the thin flow layer for alignment, after which the two-layer assembly was baked to achieve undercured PDMS-PDMS bonding (Fig. 2B). The two-layer assembly was trimmed and inverted, exposing the open-faced flow layer of the device. 0.2 μL of 1.5 μM barcoded RT primers were then spotted into the lysis module of each reaction lane and allowed to dry (Fig. 2C). By spotting the primers directly into the lysis modules, we avoid subsequent alignment steps. The two-layer chip (still undercured) with dried primers was placed atop an undercured dummy layer and bonded with heat to complete crosslinking between the layers (Fig. 2D). Finally, the assembled μCB-seq device was cut from the dummy wafer and bonded onto a #1.5 glass coverslip using oxygen plasma bonding. The result of this fabrication protocol was a valve-based multilayer microfluidic device, preloaded with barcoded RT primers at addressable locations (Fig. 2E, *Materials and Methods*).

**Fig2:**
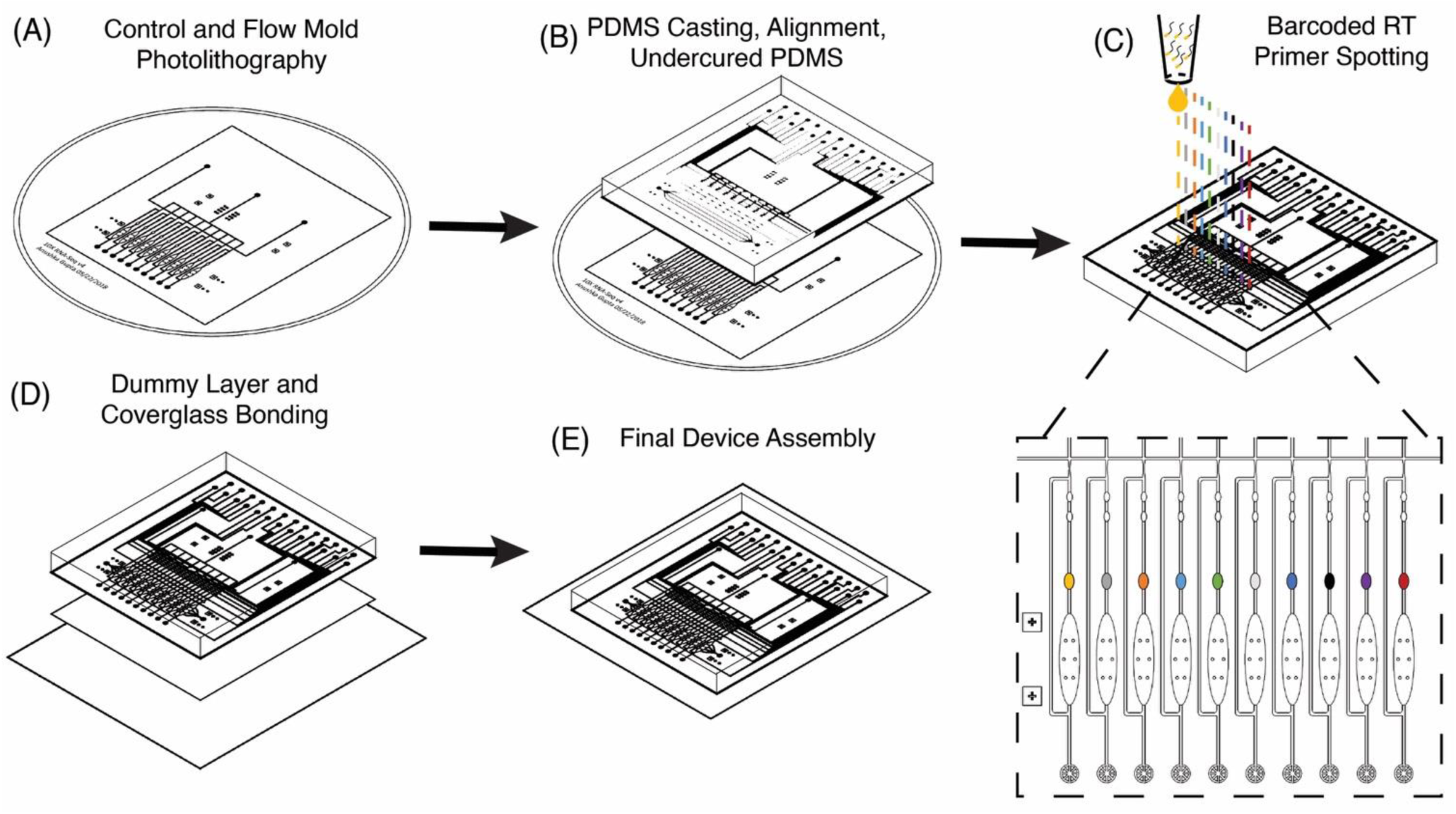
Fabrication of μCB-seq devices with barcoded RT primer spotting. (A) Photolithographic patterning of control and flow molds on Si wafers. (B) Diagram of PDMS casting and undercured PDMS bonding between the control and flow layers. (C) Detailed diagram of barcoded RT primer spotting. Unique primers are delivered to each lysis module and dried before the device is closed (D) by bonding to a PDMS dummy layer. (E) PDMS devices are then plasma bonded to a coverglass for final assembly.

### 3. μCB-seq yields high-quality scRNA-seq libraries

μCB-seq was designed to be compatible with most single-cell library preparation protocols. In this demonstration of μCB-seq, single-cell cDNA libraries were prepared by implementing the highly sensitive mcSCRB-seq protocol within the microfluidic device. mcSCRB-seq is a multiplexed 3’ counting method using cell barcodes and UMIs to acquire an absolute transcript count from each cell.^22^ We first evaluated the effectiveness of μCB-seq by generating cDNA libraries from 20 replicates of 10 pg total RNA isolated from HEK293T cells. Total RNA extracted from HEK293T cells was injected into the cell inlet and the 10 sets of isolation valves were simultaneously actuated to trap 10 pg RNA in each imaging chamber. The contents of each imaging chamber were then pushed into their respective reaction lanes for cDNA processing *(Materials and Methods)*. The cDNA libraries were then collected from the chip, pooled and prepared for high-throughput sequencing. The libraries were sequenced with Read 1 (R1) encoding the 8-nt long known barcode sequence and 10-nt long UMI and Read 2 encoding the cDNA fragment. After sequencing, all raw fastq files were analyzed using the zUMIs pipeline *(Materials and Methods)*.^33^ In zUMIs, reads with all R1 bases having quality score > 20 were mapped to the human reference genome (GrCh38) using STAR.^34^ Gene annotations were obtained from Ensembl (GRCh38.93) and filtered to remove biotypes such as pseudogenes.^35^ Quantification of aligned reads was done using the Subread package to generate expression profiles for each library.^36^ Throughout this study, genes detected were defined as those for which at least one UMI was detected. In total, all 20 libraries of purified RNA were sequenced to an average depth of 65,000 reads.

We first characterized the mapping statistics for each of the 20 total RNA libraries, which allowed us to evaluate the percentage of useful reads for downstream analysis. Across all the replicates, a median of 53% of reads mapped to exons, 11% to introns, 16% to intergenic regions, and 17% to no region in the human genome (Fig. 3A). These statistics are comparable to other 3’-barcoding-based sequencing protocols with a range of 29-57% exonic reads, 2-15% intronic reads and 6-23% unmapped reads.^37^ Detection of reads from unspliced transcripts makes μCB-seq data compatible with single-cell analyses utilizing splicing events such as RNA velocity.^38^ Here, reads mapping to the exonic regions of the genome were quantified to generate a UMI count expression matrix. These 10 pg total RNA sequencing libraries generated with μCB-seq detected a median of 3,008 unique genes with only 30,000 reads per sample (Fig. 3B). Transcript abundance was strongly correlated between μCB-seq libraries, with a median pairwise Pearson coefficient of 0.84 (n = 190 pairs) across reaction lanes and devices (Fig. 3C).

**Fig3:**
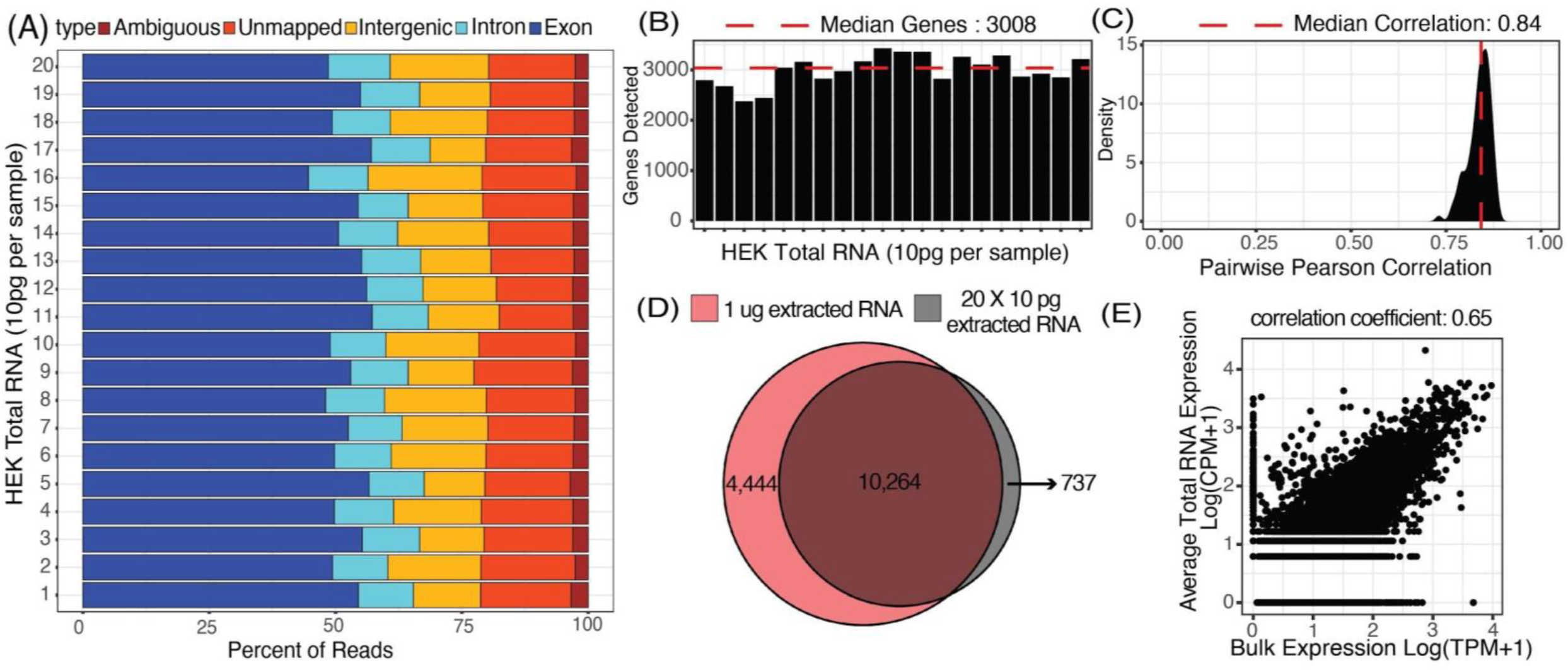
Characterization of total RNA libraries generated using μCB-Seq. 20 libraries of 10 pg total RNA extracted from HEK293T cells were sequenced using μCB-seq. (A) Distribution of percent exonic, intronic, intergenic, ambiguous and unmapped reads in each of the 20 libraries. (B) Number of genes detected (UMI count > 0) in each of the 20 libraries sequenced to a depth of 30,000 reads per sample. (C) Distribution of correlation in gene expression profile for all possible pairs of the 20 libraries (n = 190 pairs). Pearson correlation coefficients were calculated for genes detected in at least one of the 20 libraries. (D) Genes detected in a pool of the 20 libraries sequenced to a depth of ∼1.3 million reads (grey circle) compared with the genes detected in a bulk library (TPM > 0) prepared using 1µg total RNA and sequenced to the same depth (red circle). (E) Scatter plot shows correlation in gene expression profile between an average 10pg library of total RNA and the bulk library prepared using 1µg total RNA. Pearson correlation coefficient was calculated using genes detected in either bulk sample or one of the 20 total RNA libraries.

Next, we compared transcript abundance in these pseudo-single-cell libraries with typical gene expression in HEK293T cells as measured by bulk RNA-seq of HEK total RNA (1 μg, Material and Methods). For comparison, we pooled the reads from all 20 μCB-seq libraries of 10 pg total RNA for a total of 1.3 million reads and compared the genes detected against those present in 1.3 million bulk sample reads (TPM > 0). With the same total number of reads, ∼70% of genes that were present in bulk RNA-seq library of 1 μg total RNA were also detected in pooled μCB-seq libraries consisting of 200 pg RNA in total (Fig. 3D). There were over 700 genes that were detected in μCB-seq but not in bulk RNA-seq. These are likely a combination of low-abundance transcripts and transcripts that are not primed or reverse-transcribed in bulk due to molecular differences in the protocols. Transcript abundance in an average 10 pg total RNA library (averaged counts per million over all 20 replicates) correlated well with the bulk measurement (Pearson correlation = 0.65, p-value < .05, Fig. 3E). This demonstrates that μCB-seq can recapitulate expected gene expression profiles with low quantities of mRNA.

### 4. μCB-seq offers improved gene detection sensitivity

The sensitivity of a scRNA-seq protocol can be understood as the efficiency of mRNA capture and conversion into sequenceable cDNA molecules. More practically, the number of genes detected from a single cell is commonly used as a proxy for sensitivity. Gene detection sensitivity can be reduced by many sources of inefficiency, including adsorption of molecules to reaction chamber walls, inefficient reverse transcription, and transcript loss during bead cleanup steps. When molecules are lost after PCR, the information content of the library is not reduced significantly, since each transcript has many duplicates in the pool that contain the same information. Transcript loss before PCR, however, reduces the overall library complexity and severely reduces the sensitivity of the protocol. Multiplexed plate-based scRNA-seq protocols often rely on lossy bead-based cleanup to pool and concentrate single-cell cDNA libraries after RT but before PCR, a process which necessarily loses unique cDNA molecules during bead binding and elution.^22,39,40^ This loss of molecules before PCR reduces the sensitivity and gene detection capability of multiplexed scRNA-seq protocols compared to their theoretical maximum. Here, we show that microfluidic library preparation allows us to improve performance of a highly sensitive protocol by eliminating post-RT bead-based pooling altogether, because cDNA only occupies nanoliter-scale volumes on-chip.

We evaluated the sensitivity of scRNA-seq on the μCB-seq platform by sequencing the transcriptomes of single HEK cells and comparing the genes detected to single HEK cell libraries generated by mcSCRB-seq in a standard 0.3 mL 96-well plate (also described as in-tube, *Materials and Methods*). We prepared scRNA-seq libraries from 18 single cells on μCB-seq devices and 16 single cells using mcSCRB-seq in-tube. All libraries were sequenced to an average depth of 500,000 total reads per cell and downsampled to evaluate gene detection as a function of sequencing depth. The zUMIs pipeline was used to generate the count matrix for all sequencing depths, which included only exonic reads. μCB-seq consistently detected more genes and UMIs (Fig. S3), with significantly higher genes for depths >= 40,000 reads per cell (p-value < 0.01, two-group Mann-Whitney U-test, Fig. 4A). Moreover, μCB-seq libraries had a median of 21% intronic reads as compared to 15% in mcSCRB-seq (Fig. S4) which were not counted during transcript quantification, making Fig. 4A a conservative estimate of the sensitivity improvements offered by the microfluidic protocol (Fig. S5).

**Fig. 4.**
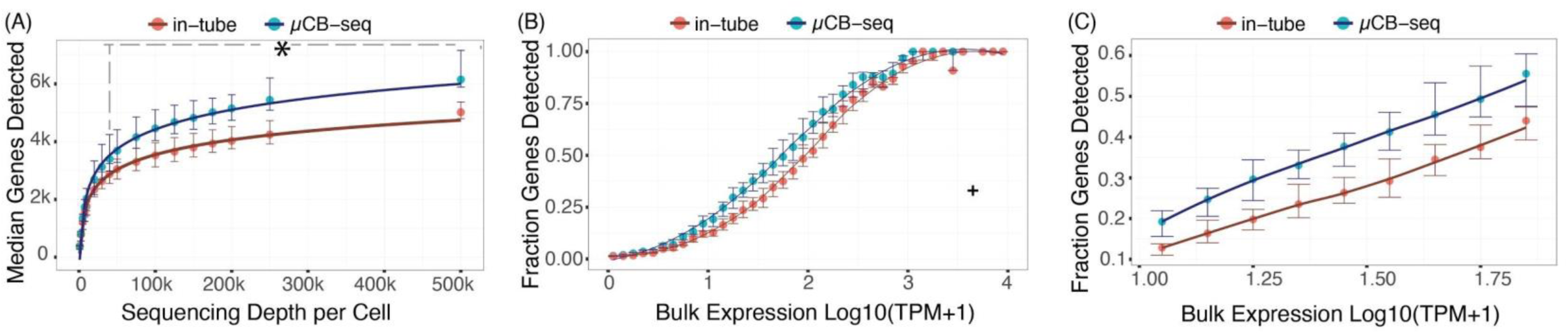
μCB-seq is more sensitive than in-tube mcSCRB-seq protocol. (A) Median genes detected for downsampled read depth across single HEK cells sequenced using μCB-seq and mcSCRB-seq. μCB-seq detected significantly higher genes for read depth >= 40,000 as tested by two-group Mann-Whitney U-test (p-value < .01). Error bars indicate the interquartile range. (B) The ratio of genes detected (UMI count > 0) in the single-cell libraries sequenced to an average depth of 200,000 reads to the genes detected in the bulk library (TPM > 0) binned by expression level (bin width = 0.1). Bulk library was prepared using 1µg total RNA and sequenced to a depth of 63 million reads. Error bars indicate interquartile range (n = 16 cells for each protocol). For a single bin (marked by +), only one out of three genes were detected in all single cells across both protocols and was considered an outlier. A Loess regression was used as a guide to the eye for this plot. (C) A magnified plot of panel (B) comparing the fraction of genes detected in the two protocols with low- and medium-abundance in bulk measurement (9 < Bulk TPM < 79).

We further evaluated the sensitivity of μCB-seq and mcSCRB-seq in-tube by comparing gene detection efficiency as a function of transcript abundance across all expression levels. Detection efficiency was calculated as the fraction of genes detected in bulk that were also detected in a single cell for a given abundance bin. Bulk library was prepared using 1μg total RNA extracted from HEK293T cells and sequenced to a depth of 63 million reads *(Materials and Methods)*. We downsampled all μCB-seq and mcSCRB-seq libraries to 200,000 reads per cell with 16 cells in each protocol. μCB-seq detected more genes than mcSCRB-seq across all expression levels, with a substantial increase in our ability to detect low- and medium-abundance transcripts (Fig. 4B and 4C).

Next, we assessed measurement precision in the μCB-seq protocol as compared to mcSCRB-seq in-tube. Variation in gene count measurements between single-cell cDNA library preparations is caused by technical variation such as pipetting, human handling errors, and sampling statistics, as well as true biological variation between cells. With microfluidics, it is possible to minimize the technical noise by automating and parallelizing library preparation reactions in lithographically defined volumes.^23,41^ As the noise associated with technical artifacts decreases, we gain statistical power to parse out real biological variation. To quantify this, we calculated the coefficient of variation (CV) for common genes detected across bulk RNA-seq, μCB-seq, and mcSCRB-seq libraries as a function of bulk expression levels. We observed slightly lower variation in μCB-seq compared to mcSCRB-seq across the entire range of bulk expression except for very highly abundant genes (TPM >= 560, Fig. S6). These results indicate that μCB-seq offers improved gene detection sensitivity with comparable measurement precision by eliminating lossy post-RT bead-based cleanup and carrying out library preparation in lithographically defined nanoliter-scale volumes.

### 5. μCB-seq links high-resolution optical images with the transcriptome of the same single cell

μCB-seq enables the collection of both imaging and sequencing data from single cells by associating known barcodes with microfluidic lane indices. As a proof-of-concept demonstration of μCB-seq, we captured high-resolution confocal images and sequenced the transcriptomes of single cells from a population of two differentially labeled cell types. We stained HEK293T cells and adipocyte precursor cells (preadipocytes)^42^ with CellBrite Green and Red cytoplasmic membrane dyes respectively *(Materials and Methods)*. The cells were then suspended and processed in three μCB-seq devices. One device processed a mix of both HEKs (n=4) and preadipocytes (n=3). The other two devices processed just HEKs (n=7), or just preadipocytes (n=6) separately. Fluorescence confocal imaging was performed while cells were isolated in the imaging chambers using 488 nm and 633 nm lasers and with a 63X magnification 0.7 NA air objective *(Materials and Methods).* The cells were then ejected into their respective reaction lanes for library preparation on-chip followed by pooled PCR. All 20 libraries were sequenced to a minimum sequencing depth of 125,000 reads per cell. After sequencing, we demultiplexed reads based on their cell barcodes, which allowed us to assign each cDNA read to the lane index and thus to the image of the cell from which the molecule originated. In this analysis, both intronic and exonic reads were used for generating a count matrix to utilize the introns detected by μCB-seq.

Fig. 5A displays representative scanning-transmission and scanning-confocal images of HEKs and preadipocytes in both green and red channels confirming differential labeling of the two cell types (Fig. S8). μCB-seq allows for high NA, and therefore high-resolution imaging, which provides the potential to draw connections between subcellular features and gene expression. Using distinct stains allowed us to determine the cell type of each captured cell prior to sequencing-based analysis. As expected, quantification of the fluorescence signal in the green and red channels completely separated the two cell-types along those two axes (Fig. 5B, *Materials and Methods*). Groups of HEKs and preadipocytes identified using image analysis also presented as two distinct cell populations upon unsupervised clustering in the principal component space (Fig. 5C, *Materials and Methods*). No technical artifacts associated with the three different devices were observed in the reduced space (Fig. S7). In this case, μCB-seq optical imaging serves as a ground truth for naïve clustering of transcriptomic data from the same cells.

**Fig. 5.**
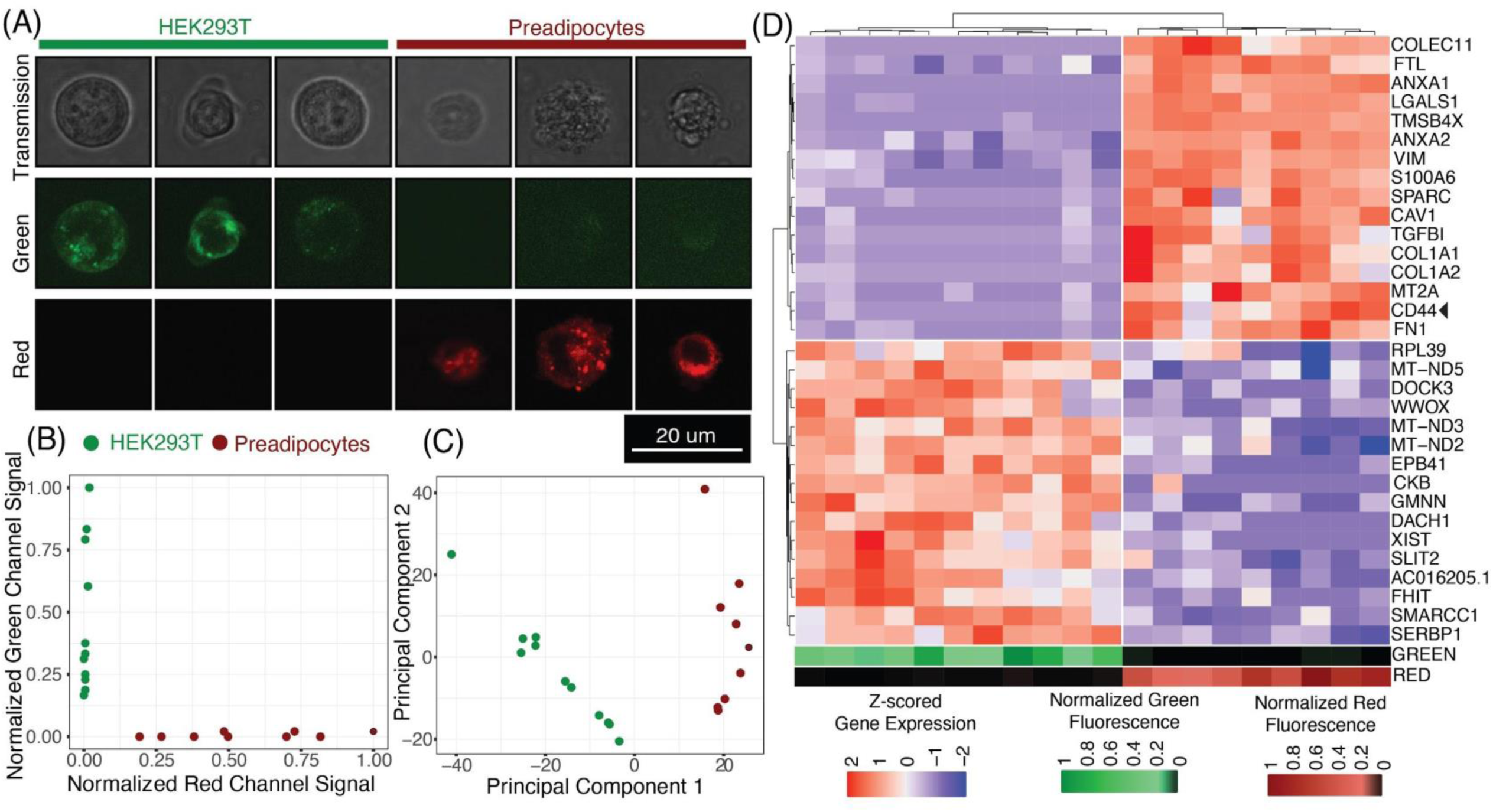
Linked imaging and sequencing using μCB-seq. (A) Montage of representative images of HEKs and Preadipocytes acquired using scanning transmission and scanning confocal microscopy in the green and red channels. HEKs and Preadipocytes were stained with CellBrite Green and Red cytoplasmic membrane dye respectively. (B) Normalized fluorescence signal in the green and red channel confocal images of both HEKs and Preadipocytes. Analysis of images for cell-mask generation and quantification of fluorescent intensities is explained in the *Methods* section. (C) Accurate identification of HEKs and Preadipocytes as two cell populations using unsupervised hierarchical clustering in the principal component space. Top 2000 most variable features were used as an input for determining the first two principal components. (D) Unsupervised hierarchical clustering using scaled expression values of top-16 upregulated genes in HEKs and Preadipocytes. Heat map shows z-scored expression values for the 32 genes. On the bottom are heat map visualizations of normalized fluorescence intensities plotted in panel (B). The heat maps for the green and red channels are ordered to accurately reflect a one-on-one correspondence between imaging and sequencing data points.

We further analyzed the sequencing dataset to understand the transcriptomic variations in this heterogeneous group of 20 cells. Differential gene expression analysis revealed 103 genes with logFC > .5 and adjusted p-value < .05 *(Materials and Methods)*. Interestingly, preadipocytes had an enriched expression of CD44, a mesenchymal stem cell surface marker which has been suggested to be expressed in adipogenic cells.^43,44^ We also performed unsupervised hierarchical clustering on the expression levels of the top 16 upregulated genes in each of the two cell types. All twenty cells were sorted into two distinct groups that accurately reflected their known cell type (Fig. 5D). These data demonstrate that μCB-seq can successfully pair high-sensitivity gene expression profiles with high-resolution fluorescence images from single cells.

## Conclusion

Microfluidic technologies have been at the core of the recent exponential increase in the throughput of scRNA-seq techniques, paving the way for undertakings such as the Human Cell Atlas Project. ^17^ However, because scRNA-seq can only record information encoded as a sequence of nucleotides, orthogonal measurements enabled by quantitative live-cell imaging, such as fluorescence staining, subcellular lipid quantification ^45^ or organelle-level pH measurements,^46^ will play an important role in the generation of a comprehensive human cell atlas. In this report, we present μCB-seq, a scalable microfluidic platform which allows us to acquire high-resolution images and generate RNA-sequencing libraries from the same single cells. μCB-seq links optical and genomic measurements with known barcodes, which are delivered to addressable locations on-chip and recovered with high efficiency during device operation, even after fabrication at 80 °C. The microfluidic device also features a modular design that allows for other multistep scRNA-seq library preparation protocols on-chip. μCB-seq’s ability to correlate optical measurements with gene expression on the single-cell level has the potential to provide insight into the relationship between genome regulation and cellular phenotypes. While this scRNA-seq demonstration uses a single barcoding step, we believe our μCB-seq barcoding approach may prove useful for many-step reactions in which aqueous samples can be automatically directed to multiple preloaded chambers for combinatorial spatial barcoding,^47^ targeted gene expression,^12^ or CRISPR-based gene editing.^48^

By using a microfluidic approach in μCB-seq for library preparation, we have eliminated post-RT bead-based cleanup, minimized operational errors, and achieved nanoliter-scale, reproducible reaction volumes. Our microfluidic approach offers improvements in sensitivity, as demonstrated by an increased gene detection efficiency. Using μCB-seq, we were also able to effectively reconstruct a large portion of the bulk transcriptome by sequencing 200 pg total RNA to a total depth of ∼1.3 million reads. The integration of on-chip valves in the device allowed us to actively select cells of interest, making the μCB-seq platform applicable for studies focusing on rare cell populations.^49^ On-chip isolation valves prevent cellular motion due to fluid flow, thereby allowing the acquisition of even prolonged spectroscopic measurements^50^ on our device. For future experiments, the throughput of μCB-seq can be increased tenfold with the current barcode list by using a microfluidic multiplexing strategy with a minimal increase in the peripheral operating equipment.^51,52^ Additionally, high-precision, low volume array spotters can be used to automate barcode preloading and increase device throughput. However, the throughput of linked imaging and sequencing measurements by μCB-seq is ultimately limited by imaging time. We believe the μCB-seq platform will be a powerful tool for investigations aiming to understand the association between a phenotype and the transcriptome, thereby gaining a high-resolution fingerprint for a particular cell population identified using other higher-throughput scRNA-seq protocols.

## Materials and Methods

### HEK293T Cell Culture and Single-Cell Suspension Preparation

HEK293T cells were obtained from the UCSF cell repository, and cultured in DMEM medium (Gibco, 10566-016) supplemented with 10% vol/vol FBS and containing 1% vol/vol Penicillin-Streptomycin (Gibco). The cell culture was maintained at 37 °C in a humidified incubator containing 5% vol/vol CO_2_. Confluent cells were passaged using TrypLE (Gibco,12563011) with a 1:25 split in a new T25 flask (Falcon, 353109). For generating HEK293T single-cell suspensions for μCB-seq vs mcSCRB-seq comparisons (Fig. 4), cells were first grown to 100% confluence. The cells were then resuspended in 1 mL TrypLE and 5 mL of growth media and centrifuged at 1,200 rpm for 4 min. After centrifugation, the supernatant was removed and the cell pellet was washed with 1 mL of PBS (Corning, 21-040-CV). The cells were centrifuged again and this process was repeated for a total of three PBS washes to remove cell debris. Finally, the concentration of the cell suspension was adjusted in ice-cold PBS to 700 cells/μL using a hemocytometer (Hausser Scientific). After this, the cell suspension was always stored on ice throughout the course of device operation. In most experiments, around 50 μL of the single-cell suspension was aspirated into a gel-loading pipette tip and placed into the device, although the full volume was rarely completely used, and it is possible to decrease this volume in situations where the sample is limited.

### Preadipocyte Cell Culture

Human preadipocytes were provided by our collaborators in the Tseng lab at Joslin Diabetes Center at Harvard. The cells were isolated from the deep neck region of a deidentified individual using the protocol in Xue et al. and immortalized to allow for cell culture and expansion.^42^ For culturing, Preadipocytes were grown in DMEM medium (Corning,10-017-CV) supplemented with 10% vol/vol FBS and containing 1% vol/vol Penicillin-Streptomycin (Gibco). The cell culture was maintained at 37 °C in a humidified incubator containing 5% vol/vol CO2. 80% confluent cells were passaged using 0.25% trypsin with 0.1% EDTA (Gibco; 25200-056) for a 1:3 split in a new 100mm cell culture dish (Corning).

### HEK293T and Preadipocyte Membrane Staining Protocol

HEK293T cells and Preadipocytes were stained with CellBrite™ Green (#30021) and Red (#30023) Cytoplasmic Membrane Labeling Kits respectively using manufacturer’s protocol. Briefly, cells were suspended at a density of 1,000,000 cells/mL in their respective normal growth medium. 5 μL or 10 μL of the Cell Labeling Solution was then added per 1 mL of cell suspension for HEKs and Preadipocytes respectively. Cells were then incubated for 20 minutes (HEKs) or 40-60 minutes (Preadipocytes) in a humidified incubator containing 5% vol/vol CO_2_. Cells were then pelleted by centrifugation at 1,200 rpm for 4 min. After centrifugation, the supernatant was removed, and cells were washed in warm (37 °C) medium. Cells were centrifuged again, and the process was repeated for a total of 3 growth medium washes for HEKs and 1-3 growth medium washes for Preadipocytes. Cells were then centrifuged a final time at 1,200 rpm for 4 minutes and resuspended in ice-cold PBS (Corning, 21-040-CV) for a final concentration of 700 cells/μL adjusted using a hemocytometer (Hausser Scientific). The cells were then stored on ice throughout the μCB-seq device operation.

### Bulk RNA-sequencing and Data Analysis

Total RNA was extracted from HEK293T cells using the RNeasy Mini Kit from Qiagen (74104) with the QIAshredder (79654) for homogenization. RNA library preparation was performed with 1ug of total RNA input quantified by Qubit fluorometer using the NEBNext Poly(A) mRNA Magnetic Isolation Module (E7335S) followed by NEBNext Ultra II RNA Library Prep Kit for Illumina (E7770S). Paired-end 2 × 150 bp sequencing for the bulk library was performed on the Illumina Novaseq platform for a coverage of approximately 63 million read pairs.

For analysing the dataset, adapters were first trimmed using trimmomatic^53^ (v0.36; ILLUMINACLIP:adapters-PE.fa:2:30:10 LEADING:3 TRAILING:3 SLIDINGWINDOW:4:15 MINLEN:36, where adapters-PE.fa is:

>PrefixPE/1

TACACTCTTTCCCTACACGACGCTCTTCCGATCT

>PrefixPE/2

GTGACTGGAGTTCAGACGTGTGCTCTTCCGATCT)

After trimming, reads were then aligned to the GRCh38 index generated using STAR. We provided the GTF file that is recommended for the 10X CellRanger pipeline as an input in STAR while generating the index. Paired-reads aligning to the exonic regions were then quantified using the featurecounts command in the Subread package. Chimeric reads and primary hits of multi-mapping reads were also counted towards gene expression levels. The same GTF file as in STAR was used as the input for transcript quantification. The fragment-counts matrix so obtained was converted to Transcripts Per Kilobase Million mapped reads (TPM) using the lengths for each gene as calculated by the featurecounts command in the Subread package. For analysis in Figure 3, reads were subsampled to a depth of 1.3 million reads using the Seqtk package (v1.3).^54^ These subsampled reads were then analysed in the exact same fashion as described above.

### Confocal imaging of HEK293Ts and Preadipocytes

Fluorescence confocal imaging of cells was performed in the imaging chamber of the μCB-seq device using an inverted scanning confocal microscope (Leica, Germany), and with a 63X 0.7 NA long-working-distance air objective. As outlined before, HEKs were stained using CellBrite™ Green dye and Preadipocytes were stained using CellBrite™ Red dye. Each cell was excited by two continuous-wave lasers, a 488 nm Ar/Kr laser and a 633 nm He/Ne laser, for concurrent imaging in the green and red channels respectively. Bandpass filters captured backscattered light from 490-590 nm at the photomultiplier tube in the green channel (Green-PMT), and from 660-732 nm at the photomultiplier tube in the red channel (Red-PMT), with the pinhole set to 1 Airy unit. A third PMT simultaneously captured a scanning transmission image using the unfiltered forward-scattered light. The imaging resolution was Rayleigh-limited, with a scanning zoom of 2.2X to achieve a Nyquist sampling rate of 207 nm per pixel (as calculated for the Ar/Kr laser with a smaller wavelength). Each image was 8-bit, grayscale and 512 × 512 pixels in size. Since individual HEK cells and Preadipocytes internalized varying amounts of membrane stain, the PMT gain which utilized the entire range of bit-depth (0-255) differed from one cell to another. Therefore, stained HEK and Preadipocyte cell suspensions were first imaged on a #1.5 coverslip for adjusting the range of Green-PMT gain (range: 524.6) and Red-PMT gain (range: 512-582). We measured a maximum gain of 524.6 in the green channel and 582 in the red channel to observe cellular features, and therefore set the background PMT gain to an even higher value of 600, to validate that lack of features in background images was not because of low PMT gain. In all our images, the focal plane was positioned at the cross-section with maximum fluorescence intensity. The final images were Kalman-integrated over 6 frames to remove noise. Images in Fig. 5A have been adjusted to highlight cellular features. However, no adjustment was done for quantitative image processing.

### Quantitative Image Processing

To quantify the fluorescence signal intensity in individual HEKs and Preadipocytes labeled using the CellBrite™ Green and Red dye respectively, we wrote a custom image analysis script in Python (v3.7.1) using the skimage package (v0.20.2) and multi-dimensional image processing (ndimage) package from the SciPy (v1.2.1) ecosystem. As explained in the confocal imaging section above, each cell had two fluorescence images, one green-channel confocal image, and one red-channel confocal image. Depending on the cell-type, one of the channels exhibited cellular signal (green for HEK and red for Preadipocytes) and the second channel conversely was a control image. For images of individual HEK cells and Preadipocytes, all green-channel and red-channel images respectively were analyzed to generate a cell mask (as detailed below). The pixels constituting the cell mask were designated as foreground pixels and the remaining pixels were designated as background pixels. The fluorescence signal to noise ratio (SNR) was then quantified as the ratio of mean foreground pixel intensity over mean background pixel intensity. The same pixel annotation (for foreground and background pixels) was also used in the control images to quantify SNR in the second channel. In essence, we quantified the SNR in both green and red channels for each cell and these values were normalized to linearly scale between 0 and 1 for Fig. 5B and Fig. 5C. For cell mask generation, grayscale images were first gaussian filtered to remove noise using the ndimage.gaussian_filter command with sigma set as 1. The filtered images were converted into binary images using Otsu Thresholding from the skimage package. Pixels with value 1 in the binarized images were annotated as foreground and pixels with value 0 were annotated as background (Fig. S9).

### Principal Component Analysis, Clustering and Differential Gene Expression Analysis

For membrane-stained HEKs and Preadipocytes, principal component analysis (PCA), clustering, and differential gene expression analysis were performed using the Seurat package (v3.1.1)^55^ in the R programming language (v3.5.2). First, the umi-count matrix generated using zUMIs at a read depth of 125,000 per cell was read using the readRDS command. The count matrix was then used to create a Seurat object with no filtering for either cells or genes. The umi-count matrix was log-normalized with a scaling factor of 10,000 using the NormalizeData command. The top 2,000 most variable genes in the full dataset were identified using the variance-stabilizing transformation (vst) method implemented by the FindVariableFeatures command. The normalized count matrix was then scaled and centered to generate the Z-scored matrix using the ScaleData command. The first and second principal components were then calculated based on the Z-scored expression values of the 2,000 variable genes using the RunPCA command and the reduced space visualization was plotted using the ggplot2 package (v3.1.0) in R.

For clustering using Seurat, first, a K-Nearest Neighbor graph (KNN) was constructed using the cell embeddings in the PCA space (K=5). The generated KNN graph was then used to construct a Shared Nearest Neighbor (SNN) graph by calculating the Jaccard index between every cell and its nearest neighbors using the FindNeighbors command. Using the SNN graph, the clusters were then identified using the FindClusters command with the resolution parameter set to 0.1. At this resolution, HEKs and Preadipocytes separated into two clusters as visualized in the PCA space (Fig. 5C). After clustering, differentially expressed genes (logFC > 0.5 and adjusted p-values < .05) between the two clusters were identified by fitting a negative binomial generalized linear model (negbinom test) on the raw umi-count matrix as implemented in the FindAllMarkers command. Z-scored expression values of the top 16 upregulated genes for each cell-type were then color mapped in a Heatmap plot using the ComplexHeatmap package.^56^ ComplexHeatmap was also used to perform unsupervised hierarchical clustering of single cells and genes using the euclidean distance metric and complete linkage classification method. Imaging heatmaps, with normalized green- and red-channel fluorescence signal as the data points, were also plotted using the ComplexHeatmap package.

### Control and Flow Mold Fabrication

Two molds, a control mold and a flow mold, were patterned on silicon wafers (University Wafers, #S4P01SP) with photolithography. Patterns for the control and flow molds were designed in AutoCAD (Autodesk) and printed onto 25,400 dpi photomasks (CAD/Art Services, Inc., Bandon, Oregon). The silicon wafers were first thoroughly cleaned using acetone, isopropyl alcohol, and water. The wafers were then baked at 150 °C for 10 min to dehydrate the surface. For the control mold, a 5 μm dummy layer of SU8-2005 (MicroChem) was first spin-coated at 3,000 rpm for 30 sec. The resist-coated mold was then baked at 65 °C for 1 min and 95 °C for 2 min and exposed to UV radiation with no mask for 10 sec. After exposure, the mold was again baked at 65 °C for 1 min and at 95 °C for 3 min and allowed to cool to room temperature. After dummy layer deposition, a dollop of SU8-2025 negative photoresist (MicroChem) was poured onto the control mold directly and then spun at 3,000 rpm for 30 sec, yielding a 25-μm layer. Then, the wafer was baked on a hotplate at 65 °C for 1 min and then at 95 °C for 5 min. The resist-coated wafer was exposed to a 150 mJ/cm^2^ dose of UV radiation through a negative mask (clear features and opaque background) imprinted with the control circuit using a photolithography aligner. After exposure, the wafer was again baked at 65 °C for 1 min and 95 °C for 5 min. The wafer was then submerged in SU-8 developer and gently agitated until the unexposed photoresist was removed, leaving the positive control features. Then, the wafer was carefully washed with isopropyl alcohol and blow-dried. The mold was baked at 150 °C for at least 20 min before further use.

The flow mold was fabricated using two photoresists to achieve multiple feature heights. The flow channels were fabricated using the positive photoresist AZ 40XT-11D (Integrated Micro Materials, Argyle, TX) and the taller reaction chambers were fabricated using the negative SU8-2025 photoresist. The flow mold was first spin-coated with a 5 μm dummy layer of SU8-2005 and processed the same as described for the control mold above. After dummy layer deposition, a dollop of AZ 40XT-11D positive photoresist was poured onto the flow wafer directly and then spun at 3,000 rpm for 30 s, yielding a 20 μm layer. After baking at 65 °C for 1 min and 125 °C for 6 min, the photoresist was then exposed to a 420 mJ/cm^2^ dose of UV light through a high-resolution positive mask containing the flow circuit design and developed in AZ400K developer. We then baked the mold again at 65 °C for 1 min and at 105 °C for 100 sec to reflow the positive photoresist and create rounded channels. Negative photoresist (SU8-2025) was then used for building the reaction chambers using the same protocol as described for the control mold above.

### PDMS Device Fabrication

Each layer of the multilayer μCB-seq device was bonded together by on-ratio (10:1) bonding of RTV-615 (GE Advanced Materials).^32^ The control and flow molds were exposed to chlorotrimethylsilane (Sigma-Aldrich) vapor for 30 minutes before soft lithography to facilitate PDMS releasing from the mold. After mixing and degassing of PDMS, 50 g of PDMS was cast onto each control mold and baked at 80 °C for 15 min to partially cure the PDMS slabs. Control ports were punched and flow molds were spin-coated with a PDMS layer at a speed of 2,000 rpm for 60 sec. Flow layers were partially cured at 80 °C for 5 min, after which control slabs were aligned and placed atop flow PDMS. PDMS assemblies were cured at 80 °C for a further 10 min, after which devices were peeled off of the Si wafer. Flow ports were punched, and assemblies were placed upside-down in preparation for primer spotting. In a clean hood, 0.2 μL of 1.5 μM barcoded RT primer was manually spotted in lysis chambers using a P2 pipette, with each lane receiving a unique, known barcode sequence (Table S1). Primers were allowed to dry while a PDMS dummy layer was spin-coated and partially cured on a blank, silanized Si wafer. Control+flow-layer PDMS assemblies were then placed onto the PDMS dummy layer for a 1.5 hr hard bake at 80 °C. Final devices were bonded to #1.5 glass coverslips by O_2_ plasma (PETS Inc.) and placed at 4 °C for storage.

### Microfluidic Device Operation

Microfluidic devices were attached to an Arduino-based pneumatic controller (KATARA) in preparation for running on-chip library preparation. Prior to single-cell experiments, the cell trapping line was flushed with nuclease-free water (nfH_2_O) and incubated with 0.2% (wt/wt) Pluronic F-127 (Invitrogen, P6867) for 1 hr, leaving downstream chambers containing barcoded primers empty. A single cell suspension was prepared and drawn into the cell trapping line by peristaltic pumping action of the integrated microfluidic valves. Triton Buffer was first prepared by combining 0.2 μL RNase Inhibitor (40 U/uL, Takara 2313A) and 3.8 μL 0.2% (v/v) Triton X-100 (Sigma, X-100). Lysis buffer was then prepared by mixing 1 μL 1:100 5x Phusion HF Buffer (NEB, B0518S) 2.5 μL Triton Buffer, 0.7 μL nfH_2_O, and 0.8 μL 1% (v/v) Tween 20 (Sigma, P7949) in a 0.2 mL PCR tube. Lysis buffer was aspirated into a gel-loading pipette tip, which was inserted into the reagent inlet and pressurized. The reagent tree was dead-end filled with lysis buffer, and the device was transferred to a confocal microscope (Leica) for cell trapping and imaging.

Cells were drawn along the cell input line by the peristaltic pump and manually trapped in the imaging chamber for imaging, which was carried out by the protocol described in *Confocal Imaging*. After imaging, the lane’s reagent valves were opened, allowing lysis buffer to push the trapped cell into the lysis module containing dried, uniquely barcoded RT primers. After the dead-end filling of the lysis module, primers were resuspended by pumping action of the microfluidic paddle above the lysis chamber. The microfluidic device was transferred to a thermal block for cell lysis at 72 °C for 1 min, after which the block was cooled to 4 °C. During cooling, the reagent inlet was flushed with 20 μL nuclease-free water and dried with air. Reverse transcription mix was then prepared in a 0.2 mL tube by mixing 0.8 μL 25 mM each dNTP mix (Thermo Fisher, R0181), 4 μL 5X Maxima H-Buffer (Thermo Fisher EP0751), 0.4 μL 100 μM E5V6 TSO (Table S2), 5 μL 30% PEG 8000 (Sigma Aldrich, 89510-250G-F), 6.4 μL nfH_2_O, 0.2 μL 1% Tween 20, and 0.2 μL 200 U/μL Maxima H-Reverse Transcriptase (Thermo Fisher EP0751). Reverse transcription mix was injected into the reagent inlet to dead-end fill the reagent tree. The isolation valves were then closed and reagent valves were opened to allow the RT mix to dead-end fill all lanes. Reverse-transcription was carried out for 90 min at 42 °C, with the peristaltic pump operating at 1 Hz to accelerate diffusive mixing of cell lysate, reverse transcription mix, and barcoded primers. Following reverse transcription, the chip was cooled to 4 °C and the reagent inlet was washed and dead-end filled with nuclease-free water. Barcoded cDNA was eluted in a volume of 1.7 μL per lane into gel loading pipette tips and pooled in a single PCR tube for downstream single-pot reactions.

Exonuclease digestion was carried out on the 17 μL of pooled library by adding 2 μL Exonuclease Buffer (10X) and 1 μL 20 U/μL ExoI (Thermo Fisher, EN0581), with no concentration steps required, followed by incubation at 37 °C for 20 min, 80 °C for 10 min, and cooling to 4 °C. Following exonuclease digestion, the following reagents were added to the library tube for PCR: 1.5 μL 1.25 U/μL Terra Direct Polymerase (Clontech, 639270), 37.5 μL 2X Terra Direct Buffer, 1.5 μL 10 μM SINGV6 Primer (Table S2), and 14.5 μL nfH_2_O. PCR was carried out with the following protocol: 3 min at 98 °C followed by 17 cycles of (15 sec at 98 °C, 30 sec at 65 °C, 4 min at 68 °C), followed by 10 min at 72 °C and a 4 °C hold. Post-PCR libraries were size-selected with AmPure XP beads (Beckman Coulter, A63880) using a 0.6:1 Beads:Library volume ratio. Final libraries were run through the Nextera XT tagmentation protocol (Illumina), with the PNEXTPT5 custom primer (Table S2) substituted for the P5 index primer as in mcSCRB-seq. Indexed libraries were pooled and sequenced on an Illumina MiniSeq Platform.

### mcSCRB-seq In-Tube Library Preparation

For mcSCRB-seq in-tube experiments, 96-well plates were first prepared with 10 barcoded primers and lysis buffer according to the mcSCRB-seq protocol, with the only difference being the use of μCB-seq RT primers instead of standard mcSCRB-seq ones. For single HEK cell experiments, the CellenONE X1 instrument was used to individually deliver a single HEK cell into each well. Following cell delivery, the mcSCRB-seq protocol was followed directly, but with a 1:1 ratio of AmPure XP beads to pool all cDNA after RT as opposed to the manual bead formulation from standard mcSCRB-seq.

### HEK Single-Cell and HEK Total RNA Sequencing Data Processing

Filtering, demultiplexing, alignment, and UMI/gene counting were carried out on the zUMIs pipeline for all samples, using the GRCh38 index for STAR alignment. We provided the GTF file that is recommended for the 10X CellRanger pipeline for standardization of gene counts. Reads with any barcode or UMI bases under the quality threshold of 20 were filtered out, and known barcode sequences were supplied in an external text file. UMIs within 1 hamming distance were collapsed to ensure that molecules were not double-counted due to PCR or sequencing errors. For this analysis, cell barcodes were not collapsed based on their hamming codes. For the Total RNA μCB-seq dataset (TC012), the quality of the 3rd base of Read 1 was poor due to the fact that all barcodes in the sequencing run had an Adenine at that position. Therefore, fastq files for this dataset were edited to remove the third base, and truncated barcode sequences were provided to zUMIs to match. This modification did not affect the information content or quality of the processed library.

## Data Accessibility

Yaml files for zUMIs analysis of HEK Total RNA, single HEK cells and single HEK and Preadipocyte datasets are provided in the *streetslab* GitHub repository and can be found using this link (https://github.com/streetslab/ucb-seq-processing). Downstream data tidying and analysis was carried out in a Jupyter notebook with an R kernel, which can also be found in the repository. The CAD file with μCB-seq device design can be downloaded from the same GitHub repository.

## Acknowledgements

The authors would like to thank Prof. Yu-Hua Tseng for providing adipocyte precursors. This publication was supported by the National Institute of General Medical Sciences of the National Institutes of Health under award number R35GM124916. AG is supported by the UC Berkeley Lloyd Fellowship in Bioengineering. AS is a Chan Zuckerberg Investigator.

