## Supplementary Figures for "μCB-seq: Microfluidic cell barcoding and sequencing for high-resolution imaging and sequencing of single cells"

Electronic Supplementary Information (ESI)

**Author information**

[a] University of California, Berkeley, Department of Bioengineering, Berkeley, CA, 94720, United States of America

[b] University of California, Berkeley, Department of Materials Science and Engineering, Berkeley, CA, 94720, United States of America

[c] UC Berkeley-UC San Francisco Graduate Program in Bioengineering, Berkeley, CA, 94720, United States of America

[d] Chan Zuckerberg Biohub, San Francisco, CA, 94158, United States of America

* These authors contributed equally to this work

[+] **Corresponding Author:** Aaron M. Streets, PhD

Assistant Professor, Department of Bioengineering, University of California, Berkeley, California, 94720, United States of America

**Funding**: This publication was supported by the National Institute of General Medical Sciences of the National Institutes of Health under award number R35GM124916. Anushka Gupta is supported by the UC Berkeley Lloyd fellowship in Bioengineering. AMS is a Chan Zuckerberg Investigator.

**Conflicts of interest:** There are no conflicts to declare.

#### Supplemental Note

##### Imaging Chamber Volume Measurement for Trapping 10 pg of Total RNA

When measuring chamber volume for Total RNA experiments in the μCB-seq device (Figure 3), we initially observed a difference in height between the μCB-seq flow molds and the channels of the finalized PDMS μCB-seq devices. Flow molds were measured by Dektak profilometer, giving an imaging chamber height of 29 μm. When imaging the corresponding chamber on the μCB-seq device via Coherent anti-Stokes Raman spectroscopy (CARS), we recorded a chamber height of 53.5 μm. Profilometry was not feasible for the closed μCB-seq device, so we elected to conservatively use the CARS measurement at the risk of overestimating volume and loading less than 10 pg Total RNA into the μCB-seq device. To measure chamber volume, we pressurized the isolation valves on a μCB-seq device and acquired a z-stack of the resultant air-filled imaging chamber. Images were thresholded in ImageJ and manually outlined to record the cross-sectional area of each imaging chamber slice. The volume of the chamber was estimated by a Riemann sum to ensure that chamber volume erred on the larger side. The chamber volume measured by this method was 1.88 nL, which resulted in our conservative input concentration of 5.31 ng/μL Total RNA to ensure no more than 10 pg of RNA was processed in each lane of the μCB-seq device.

### Supplemental Figures

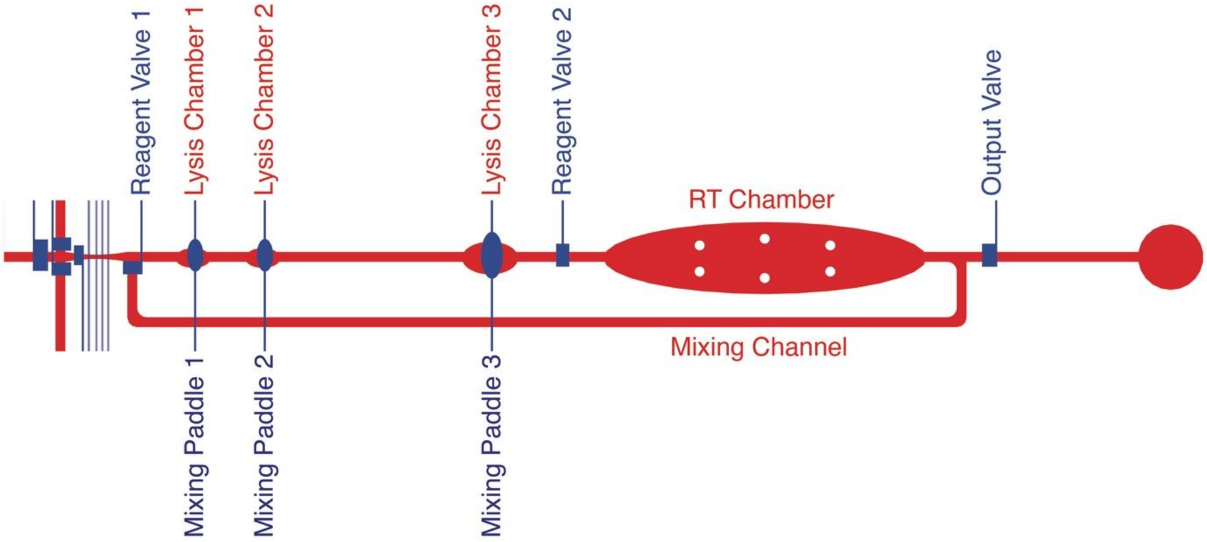


**Figure S1 Detailed schematic of a single reaction lane on the $\mu$CB-seq device.** The lysis module has 3 reaction chambers and the RT module has 1 reaction chamber connected to the mixing channel. Both lysis and RT modules are separated from each other by the two reagent valves. RT primers with known barcode sequences are spotted in the *Lysis Chamber 3* of each reaction lane. Positioned atop each of the reaction chambers in the lysis module are mixing paddles, which are actuated to resuspend barcoded RT primers in lysis buffer and circulate the relatively viscous RT mix throughout the mixing channel.

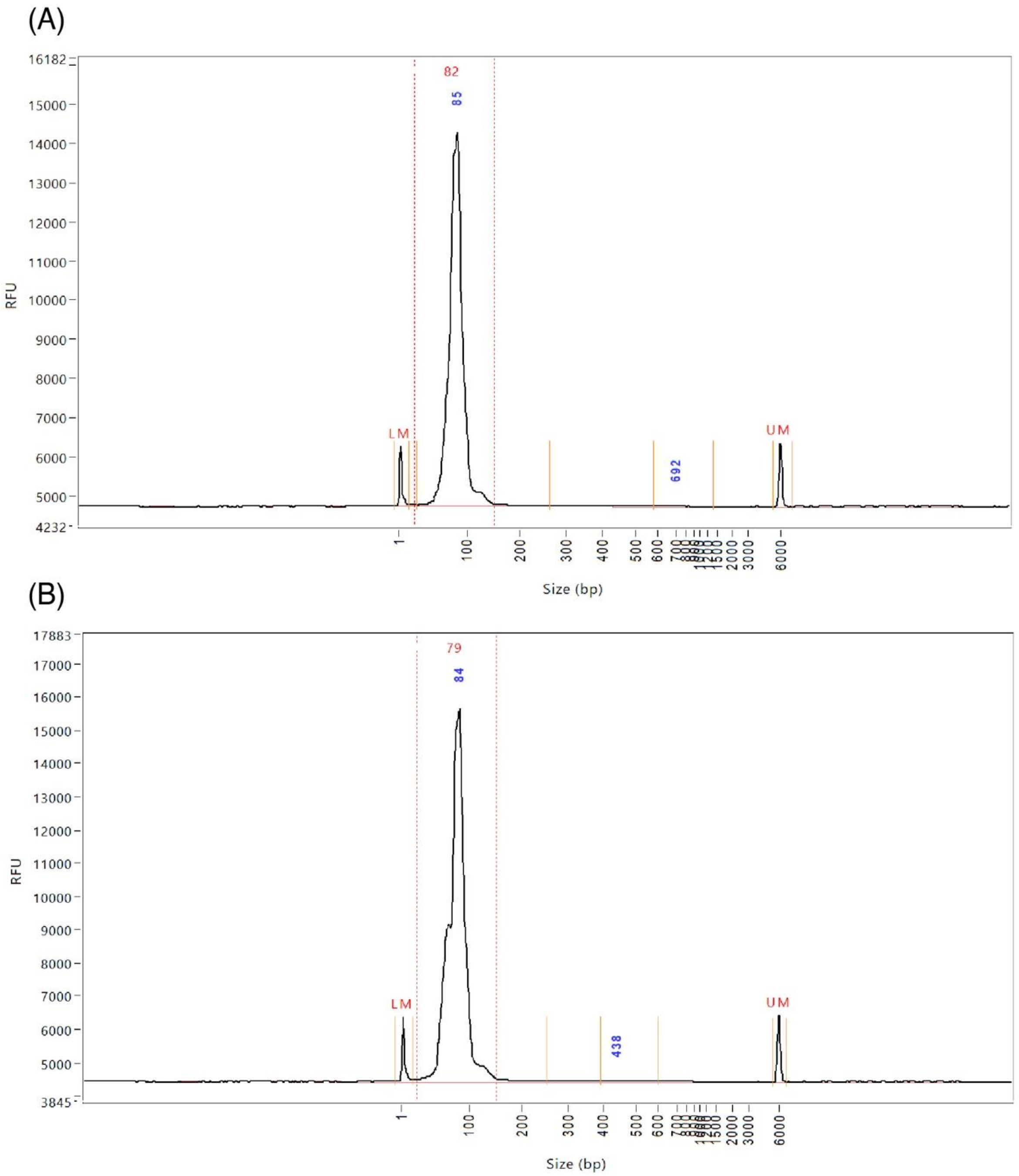


**Figure S2 Validation of intact RT primer recovery from a PDMS slab after baking.** Fragment analysis size distribution traces for barcoded primers that were suspended in nuclease-free water at RT and (A) left in the original tube or (B) spotted on PDMS, dried, baked at 80 °C and recovered by resuspending in nuclease-free water.

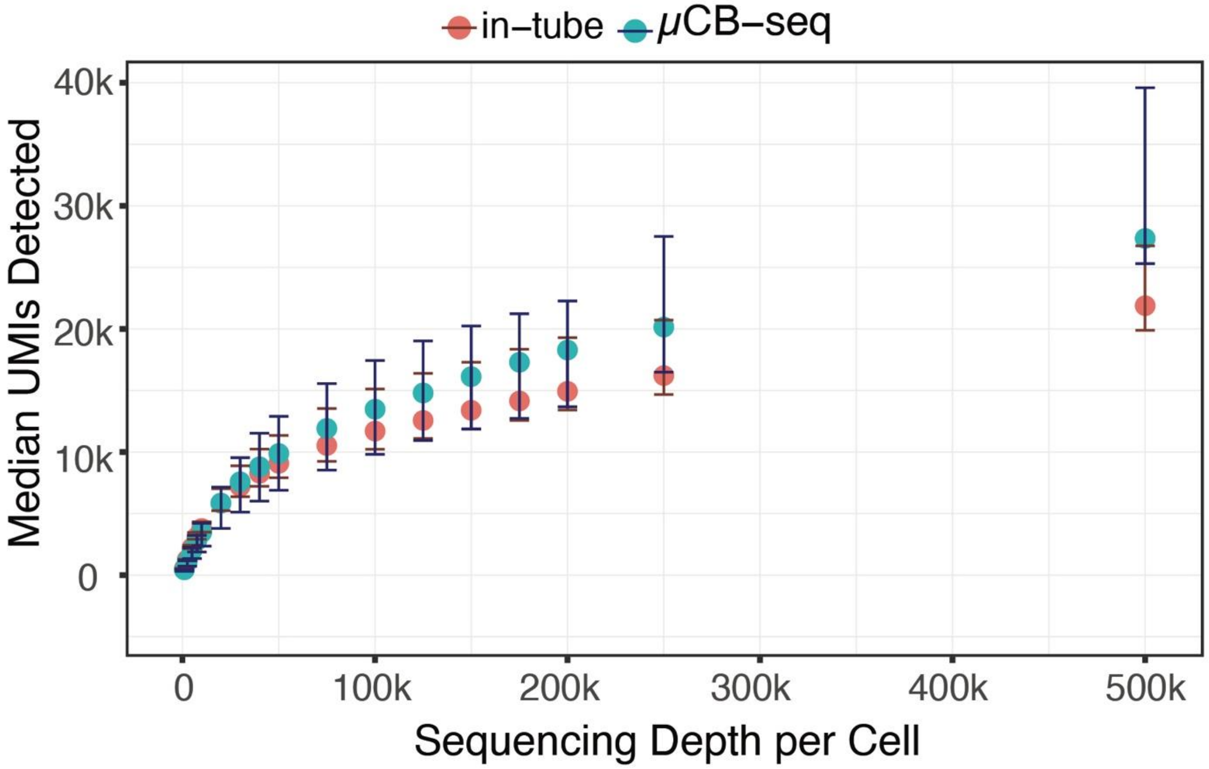


**Figure S3** Median UMIs detected for downsampled read depth across single HEK cells sequenced using $\mu$CB-seq (n = 16) and mcSCRB-seq in-tube (n = 16). Error bars indicate the interquartile range.

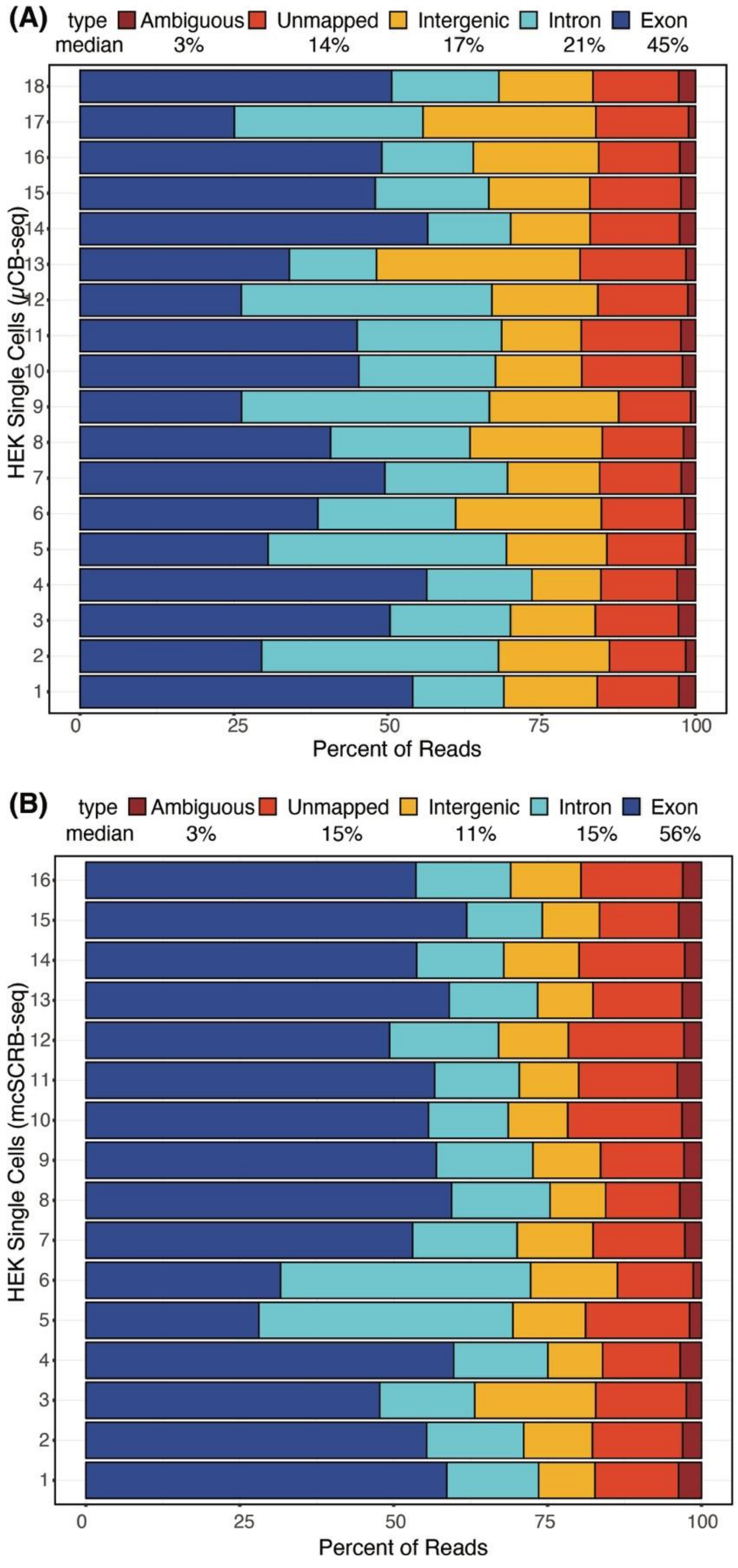


**Figure S4** Mapping Statistics for single HEK Cells sequenced using (A) μCB-seq and (B) mcSCRB-seq in-tube.

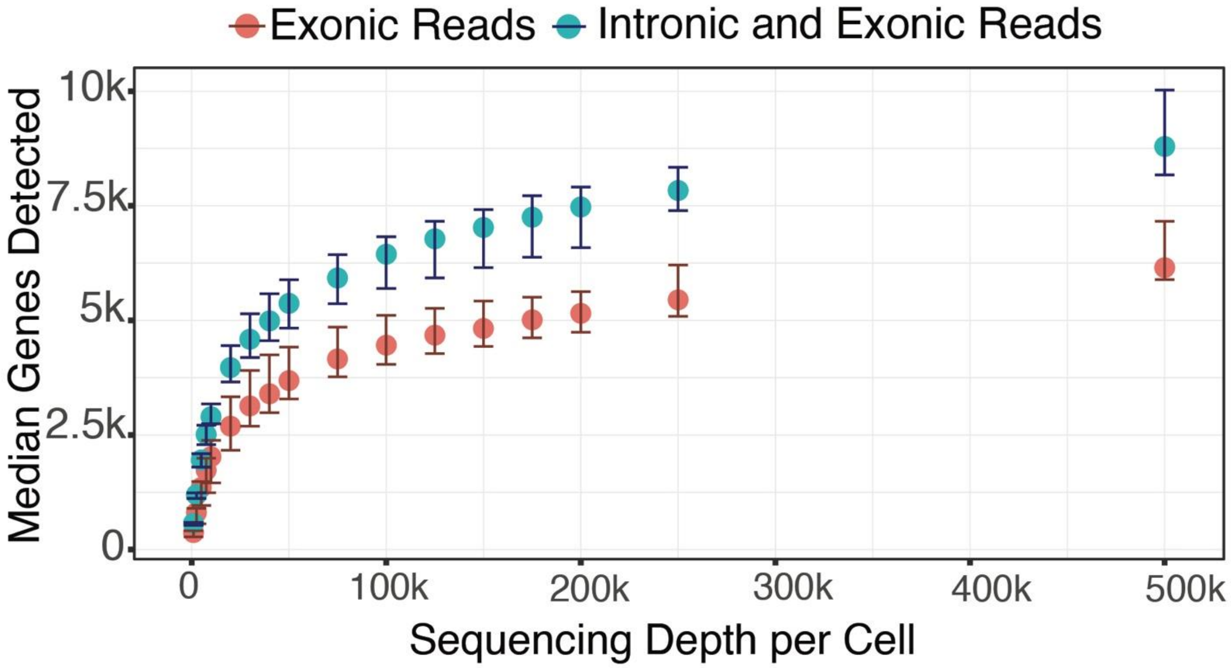


**Figure S5** Median genes detected using only exonic or both exonic and intronic reads for downsampled read depths across single HEK cells (n=16) sequenced using $\mu$CB-seq. Error bars indicate the interquartile range.

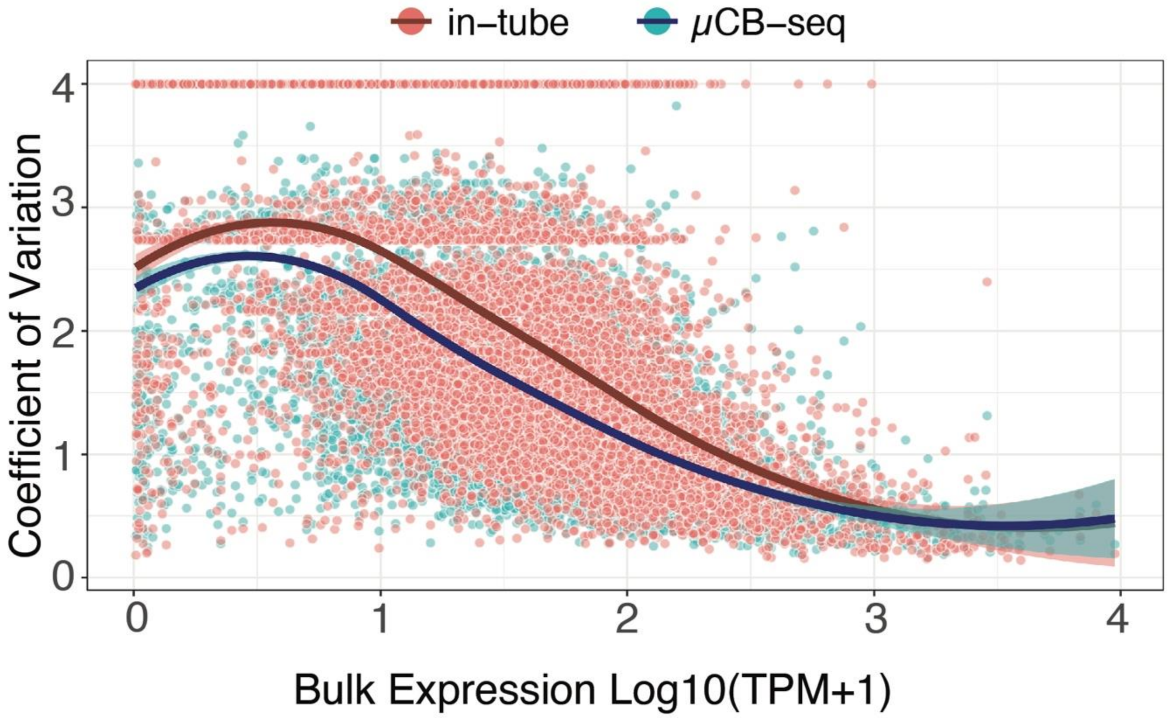


**Figure S6 $\mu$CB-seq and in-tube mcSCRB-seq protocol have comparable precision.** The coefficient of variation for each gene (SD normalized by the mean) is plotted against its bulk expression for HEK cells sequenced using $\mu$CB-seq (n=16) and mcSCRB-seq in-tube (n=16). HEK Cells were sequenced to a depth of 200,000 reads and bulk RNA-seq library was prepared using 1ug HEK total RNA sequenced to a depth of 63 million reads. CV was calculated for all common genes detected in bulk RNA-seq, $\mu$CB-seq -seq and mcSCRB-seq libraries. The highlighted region displays the 95% confidence interval around the smooth fit as determined by loess regression.

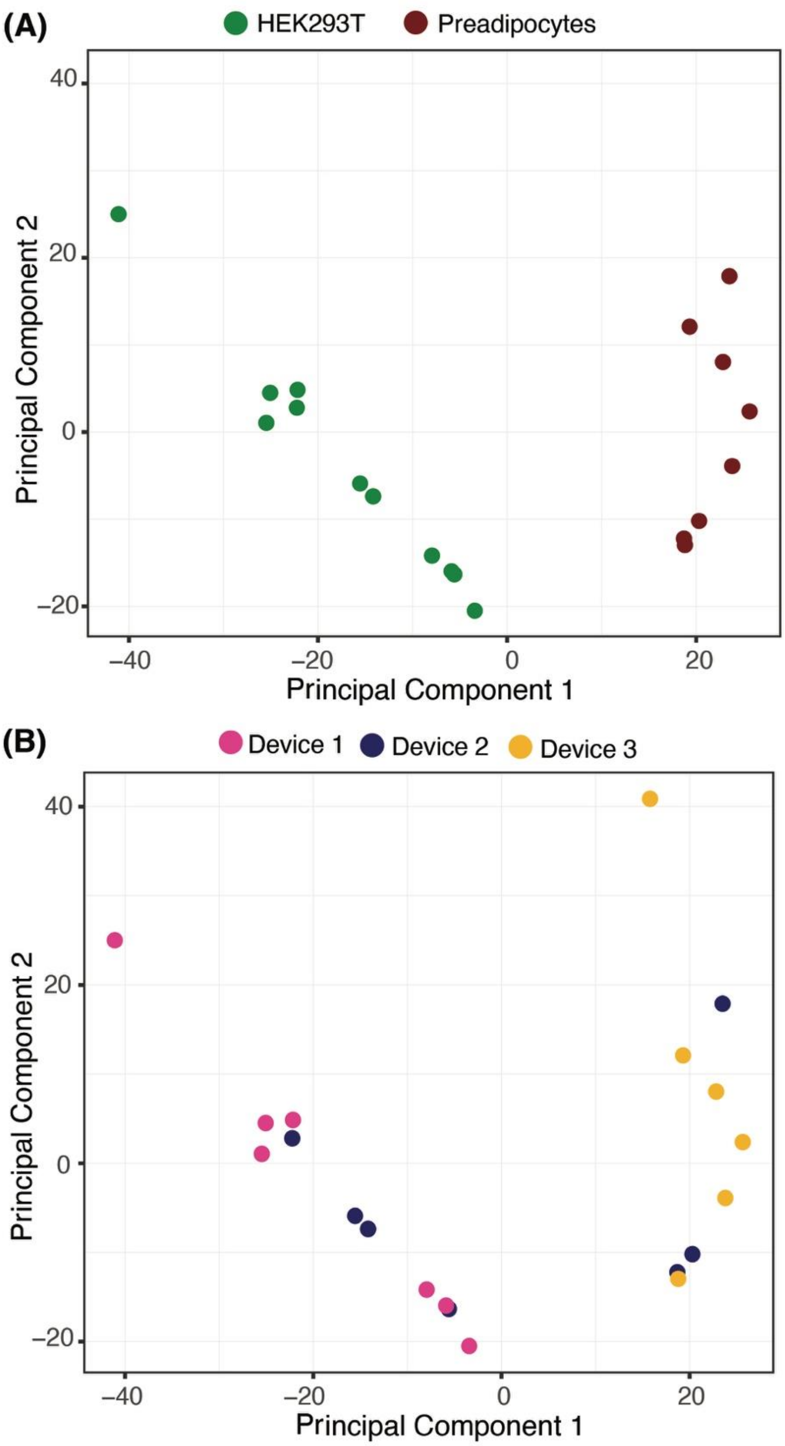


**Figure S7** Annotation of HEK293T cells and Preadipocytes in Principal Component Space based on (A) Cell-clusters identified using unsupervised hierarchical clustering in the PCA space and (B) $\mu$CB-seq devices on which cells were processed. Device 1 processed just HEKs (n=7), Device 2 processed a mix of both HEKs (n=4) and preadipocytes (n=3), and Device 3 processed just Preadipocytes (n=6)

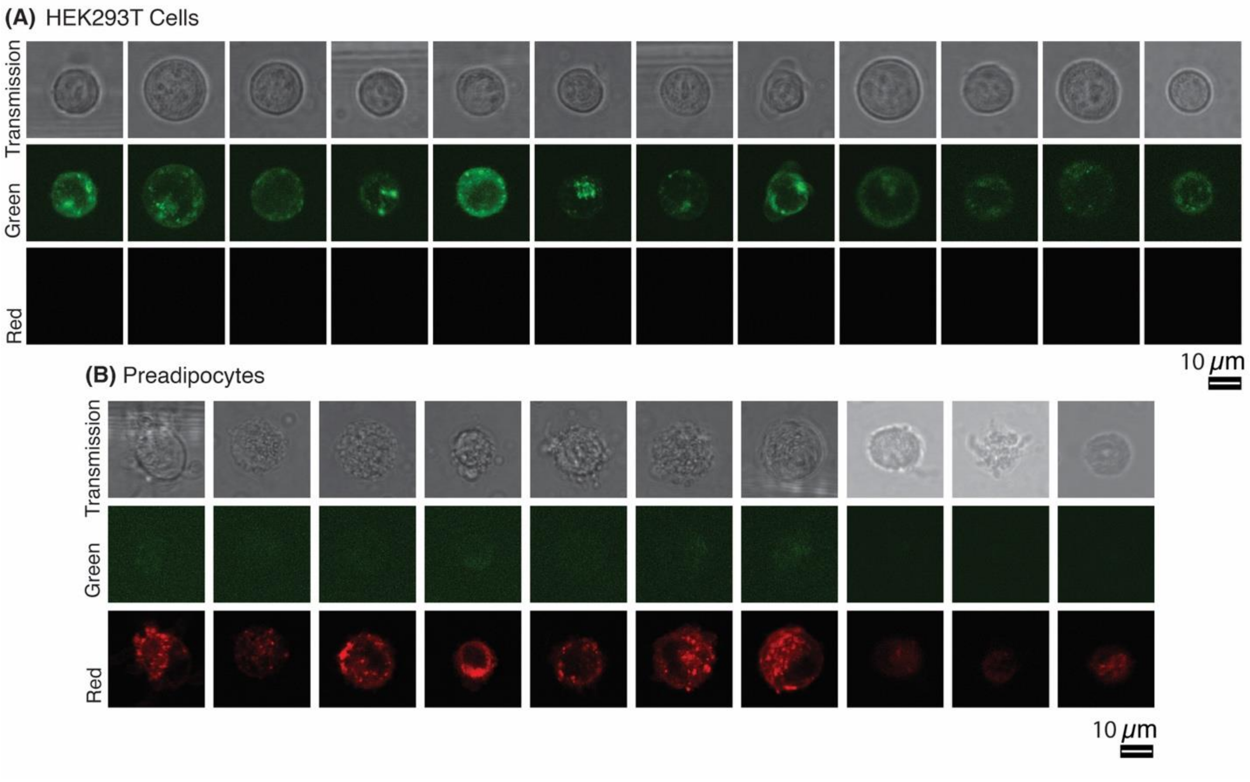


**Figure S8** Scanning transmission and two-channel fluorescent confocal images of all (A) HEK293T cells and (B) Preadipocytes stained using CellBrite™ Green and Red dye respectively.

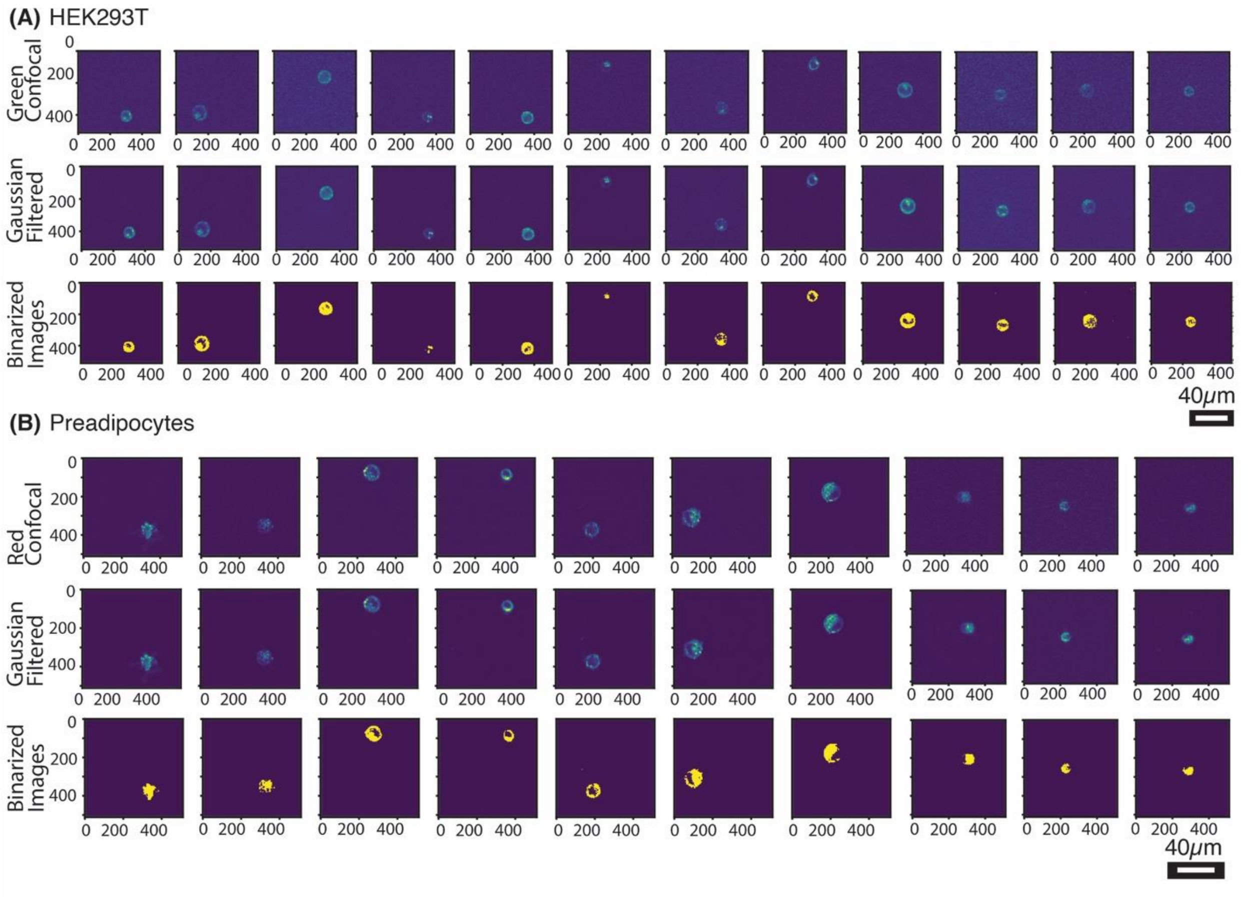


**Figure S9** Fluorescent confocal, Gaussian filtered and Otsu thresholded images of all (A) HEK293T cells and (B) Preadipocytes analyzed in Figure 5 in the main text. Detailed image analysis steps are explained in the *Materials and Methods* section.

#### Supplemental Tables

**Table S1** RT Primers with known barcode sequences used in $\mu$CB-seq. [1] Barcodes bc1-bc10 were used for experiments on HEK293T Total RNA and HEK293T single cells (Fig. 3 and Fig. 4), whereas underlined barcodes were used for the imaging and sequencing of HEK293T cells and Preadipocytes (Fig. 5). The underlined subset of ten barcodes was selected to ensure sequence diversity at every barcode base for optimal next-generation sequencing performance without PhiX spike-ins.

| Barcode Number | Barcode Sequence | Final Primer Sequence |
|---|---|---|
| <u>bc1</u> | TCACAGCA | ACACTCTTTCCCTACACGACGCTCTTCCGATCTTCACAGCA NNNNNNNNNNTTTTTTTTTTTTTTTTTTTTTTTTTTTTTTVN |
| <u>bc2</u> | GTAGCACT | ACACTCTTTCCCTACACGACGCTCTTCCGATCTGTAGCACT NNNNNNNNNNTTTTTTTTTTTTTTTTTTTTTTTTTTTTTTVN |
| <u>bc3</u> | ATAGCGTC | ACACTCTTTCCCTACACGACGCTCTTCCGATCTATAGCGTC NNNNNNNNNNTTTTTTTTTTTTTTTTTTTTTTTTTTTTTTVN |
| bc4 | CTAGCTGA | ACACTCTTTCCCTACACGACGCTCTTCCGATCTCTAGCTGA NNNNNNNNNNTTTTTTTTTTTTTTTTTTTTTTTTTTTTTTVN |
| bc5 | CTACGACA | ACACTCTTTCCCTACACGACGCTCTTCCGATCTCTACGACA NNNNNNNNNNTTTTTTTTTTTTTTTTTTTTTTTTTTTTTTVN |
| bc6 | GTACGCAT | ACACTCTTTCCCTACACGACGCTCTTCCGATCTGTACGCAT NNNNNNNNNNTTTTTTTTTTTTTTTTTTTTTTTTTTTTTTVN |
| bc7 | ACATGCGT | ACACTCTTTCCCTACACGACGCTCTTCCGATCTACATGCGT NNNNNNNNNNTTTTTTTTTTTTTTTTTTTTTTTTTTTTTTVN |
| bc8 | GCATGTAC | ACACTCTTTCCCTACACGACGCTCTTCCGATCTGCATGTAC NNNNNNNNNNTTTTTTTTTTTTTTTTTTTTTTTTTTTTTTVN |
| bc9 | ATACGTGC | ACACTCTTTCCCTACACGACGCTCTTCCGATCTATACGTGC NNNNNNNNNNTTTTTTTTTTTTTTTTTTTTTTTTTTTTTTVN |
| bc10 | GCAGTATC | ACACTCTTTCCCTACACGACGCTCTTCCGATCTGCAGTATC NNNNNNNNNNTTTTTTTTTTTTTTTTTTTTTTTTTTTTTTVN |

| | | |
|---|---|---|
| bc13 | TGCTACAG | ACACTCTTTCCCTACACGACGCTCTTCCGATCTTGCTACAG NNNNNNNNNNTTTTTTTTTTTTTTTTTTTTTTTTTTTTTTVN |
| bc15 | CGCTATGA | ACACTCTTTCCCTACACGACGCTCTTCCGATCTCGCTATGA NNNNNNNNNNTTTTTTTTTTTTTTTTTTTTTTTTTTTTTTVN |
| bc26 | ATGCACGT | ACACTCTTTCCCTACACGACGCTCTTCCGATCTATGCACGT NNNNNNNNNNTTTTTTTTTTTTTTTTTTTTTTTTTTTTTTVN |
| bc40 | TATGCACG | ACACTCTTTCCCTACACGACGCTCTTCCGATCTTATGCACG NNNNNNNNNNTTTTTTTTTTTTTTTTTTTTTTTTTTTTTTVN |
| bc47 | CATCGTGA | ACACTCTTTCCCTACACGACGCTCTTCCGATCTCATCGTGA NNNNNNNNNNTTTTTTTTTTTTTTTTTTTTTTTTTTTTTTVN |
| bc82 | CCAGTTAG | ACACTCTTTCCCTACACGACGCTCTTCCGATCTCCAGTTAG NNNNNNNNNNTTTTTTTTTTTTTTTTTTTTTTTTTTTTTTVN |
| bc92 | GGCATTGT | ACACTCTTTCCCTACACGACGCTCTTCCGATCTGGCATTGT NNNNNNNNNNTTTTTTTTTTTTTTTTTTTTTTTTTTTTTTVN |

**Table S2** Sequences of DNA primers used in both mcSCRB-seq in-tube experiments and on $\mu$CB-seq devices for off-chip library preparation. Same primer sequences as in mcSCRB-seq [2] are used in this work. /5Biosg/ indicates a 5' Biotin, *_ indicates a phosphorothioated nucleotide, and r_ indicates an RNA base.

| **Primer** | **Sequence** |
|---|---|
| SINGV6 | /5Biosg/ACACTCTTTCCCTACACGACGC |
| P5NEXTPT5 | AATGATACGGCGACCACCGAGATCTACACTCTTTCCCTACACGACGCTCTTCCG*A*T*C*T |
| E5V6 TSO | CGCACACTCTTTCCCTACACGACGCrGrGrG |
